# Signatures of adaptation at key insecticide resistance loci in *Anopheles gambiae* in Southern Ghana revealed by low-coverage WGS

**DOI:** 10.1101/2024.01.17.575856

**Authors:** Tristan P.W. Dennis, John Essandoh, Barbara K. Mable, Mafalda S. Viana, Alexander E. Yawson, David. Weetman

**Author notes:** Equal author contribution (Tristan P.W. Dennis, John Essandoh).

## Abstract

Resistance to insecticides and adaptation to a diverse range of environments present challenges to *Anopheles gambiae s.l.* mosquito control efforts in sub-Saharan Africa. Whole-genome-sequencing is often employed for identifying the genomic basis underlying adaptation in *Anopheles*, but remains expensive for large-scale surveys. Low-coverage whole-genome-sequencing (lcWGS) can identify regions of the genome involved in adaptation at a lower cost, but is currently untested in *Anopheles* mosquitoes. Here, we use lcWGS to investigate population genetic structure and identify signatures of local adaptation in *Anopheles* mosquitoes across southern Ghana. In contrast to previous analyses, we find no structuring by ecoregion, with *Anopheles coluzzii* and *Anopheles gambiae* populations largely displaying the hallmarks of large, unstructured populations. However, we find signatures of selection at insecticide resistance (IR) loci that appear ubiquitous across ecoregions in *An. coluzzii,* and strongest in forest ecoregions in *An. gambiae*. In the IR gene *Cyp9k1*, we find species-specific alleles under selection, suggesting interspecific variation in the precise mechanism of resistance conferred by *Cyp9k1*. Our study highlights resistance candidate genes in this region, and validates lcWGS, potentially to very low coverage levels, for population genomics and exploratory surveys for adaptation in *Anopheles* taxa.

## Introduction

The *Anopheles gambiae* species complex is marked by incredible genetic diversity ^1,2^ that has likely contributed to its adaptation to a diverse range of naturally-occurring and anthropogenic ecologies across sub-Saharan Africa. For example, *An. coluzzii* often breed in water sources associated with human activity (e.g. irrigation ditches, reservoirs, rice fields), and can have elevated pollution-tolerance^3–5^. By contrast, *An. gambiae* prefer humid environments, are highly anthropophilic, and tend to breed in transient, rain-dependent habitats^6^. Even within species of the *An. gambiae* complex, local genetic and ecological adaptation^7,8^ and subsequent variation in epidemiologically important traits, such as, resilience to aridity^7,9–11^, host preference and resting behaviour^12^, seasonality and preference of breeding site^6,7^ all have important implications for the design and implementation of vector surveillance and control programmes^13^. Moreover, knowledge of how environmental and geographic factors constrain gene flow can help to predict the spread of insecticide resistance (IR)^14,15^ or genetic control (e.g. gene drives) through a population^16^.

Genomic signatures of local adaptation often manifest as regions of the genome or polymorphisms displaying elevated genetic differentiation (*Fst*) among populations^17^. Whole-genome-sequencing (WGS) enables identification of *Fst* outlier regions in genome scans^18^ which, in *Anopheles spp.,* have been instrumental in identifying the genomic determinants of epidemiologically critical traits such as insecticide resistance^19^, introgression at IR loci between *Anopheles* taxa^15,20^, environmental adaptation^9,21^, and cryptic speciation^22,23^. However, broad-range surveys indicate that *Anopheles* populations, particularly those in West Africa, are exceptionally diverse and exhibit little structure over vast spatial scales^1,2^, suggesting that inference of population genetic structure over fine scales will be difficult. As such, it is possible that with sample sizes sufficiently large to capture representative allele frequencies of massive, diverse, populations, the expense of WGS may often be prohibitive. Low-coverage WGS (lcWGS) is an approach that, by reducing per-individual sequencing coverage, enables sequencing of more individuals and therefore capture of more accurate population-level allele frequencies^24,25^, while retaining individual information for many analyses that can use genotype information inferred from likelihood-based methods^26–28^ (e.g. PCA, ADMIXTURE, relatedness, inference of inbreeding). lcWGS offers promise especially for analyses relying on allele-frequency estimation - for example, in the exploratory analysis of population structure, and identification of regions of the genome under selection^29,30^ (e.g. in response to insecticide selection pressures, or local adaptation to environment) as a prelude to further investigation of specific genotypes and populations using deeper WGS. For example, lcWGS was used to identify rapid adaptation in response to fisheries-induced size selection in Atlantic silversides^31^, and environmental local adaptation at polymorphic inversions in the seaweed fly^32^. To date, however, this approach has remained untested in *Anopheles* populations.

Local adaptation and ecological divergence are often associated with transitions between biomes and environmental heterogeneity in *Anopheles gambiae*^5,21,33–35^, as well as other mosquitoes. For example in *An. funestus,* local adaptation to breeding in irrigated rice fields vs natural swamps is concentrated in chromosomal inversions^38^, as is adaptation to aridity in *An. gambiae*^9,10,21^. Insecticide pressures specific to certain habitats (e.g. land-use) may also confer habitat-specific signals of local adaptation that are related to ecology^4,5^. For example, different agricultural practices are associated with variations in the frequencies of resistance mechanisms in *An. gambiae* s.l. in Côte d’Ivoire^39^. In southern Ghana, differentiation at microsatellite loci between the four main ecozones in the region^40^: mangrove, savannah, and deciduous and rainforest ecoregions has been reported^33^, suggesting that local adaptation to ecology may be occurring in this region, though whether this is due to adaptation to the environment, or to selection pressures associated with differential pesticide use, is currently unknown.

Here, we conduct a lcWGS study of *Anopheles coluzzii* and *Anopheles gambiae* samples collected from the four main ecoregions in southern Ghana^40^ to answer the questions: 1) are there genomic signatures of adaptation to specific ecoregions? 2) Do insecticide resistance loci show signs of structuring by ecoregion? 3) And does lcWGS represent a viable option toward reduced-cost vector WGS studies?

## Results

### Population structure

A total of 314 *An. gambiae s.l.* larvae were sampled from across the four main ecoregions of southern Ghana^40^, (**Figure 1**): Rainforest (RF), Deciduous Forest (DF), Coastal Savannah (CS) and Mangrove Swamp (MS). (N.B. due to a lack of *An. gambiae* samples from MS, *An. gambiae* analyses were restricted to CS, DF and RF). Samples were whole-genome-sequenced to a target depth-of-coverage of ∼10X. Species PCR (see **Materials and Methods**) identified 159 *Anopheles coluzzii* individuals and 155 *Anopheles gambiae* individuals (**Supplementary Table 1**). Sample counts per-species and per-ecoregion are detailed in **Table 1**. To explore population structure and possible correspondence to ecoregion within the two species, we ran PCA and ADMIXTURE on *An. coluzzi* (**Figure 2A-C**) and *An. gambiae* (**Figure 2D-F**) samples. *Anopheles coluzzii* exhibited a single large cluster corresponding to the majority of the samples from all 4 ecoregions. PC1 and PC2, and PC3 and PC4, show outlier individuals and small subclusters, the most divergent of which came from the DF zone **(Figure 2A and 2B).** ADMIXTURE analysis of chromosome arm 3L GLs from *An. coluzzii* supported the presence of one major and one minor cluster, with no apparent clustering pattern by ecoregion (**Figure 2C**), and only 5 individuals predominantly belonging to Cluster 2, corresponding to the DF outroup samples in **2A** and **2B**. PCA of *An gambiae* 3L GLs showed most individuals belonging to one large cluster, with outlying subclusters again corresponding to samples from the DF zone (**Figure 2D and E**). ADMIXTURE analysis of chromosome arm 3L GLs from *An. gambiae* supported the presence of one major and two minor clusters, with no apparent clustering pattern by ecoregion. Unlike *An. coluzzii*, the minor admixture clusters did not correspond to outlying subgroups in the PCA. (**Figure 2F**)

**Figure 1:**
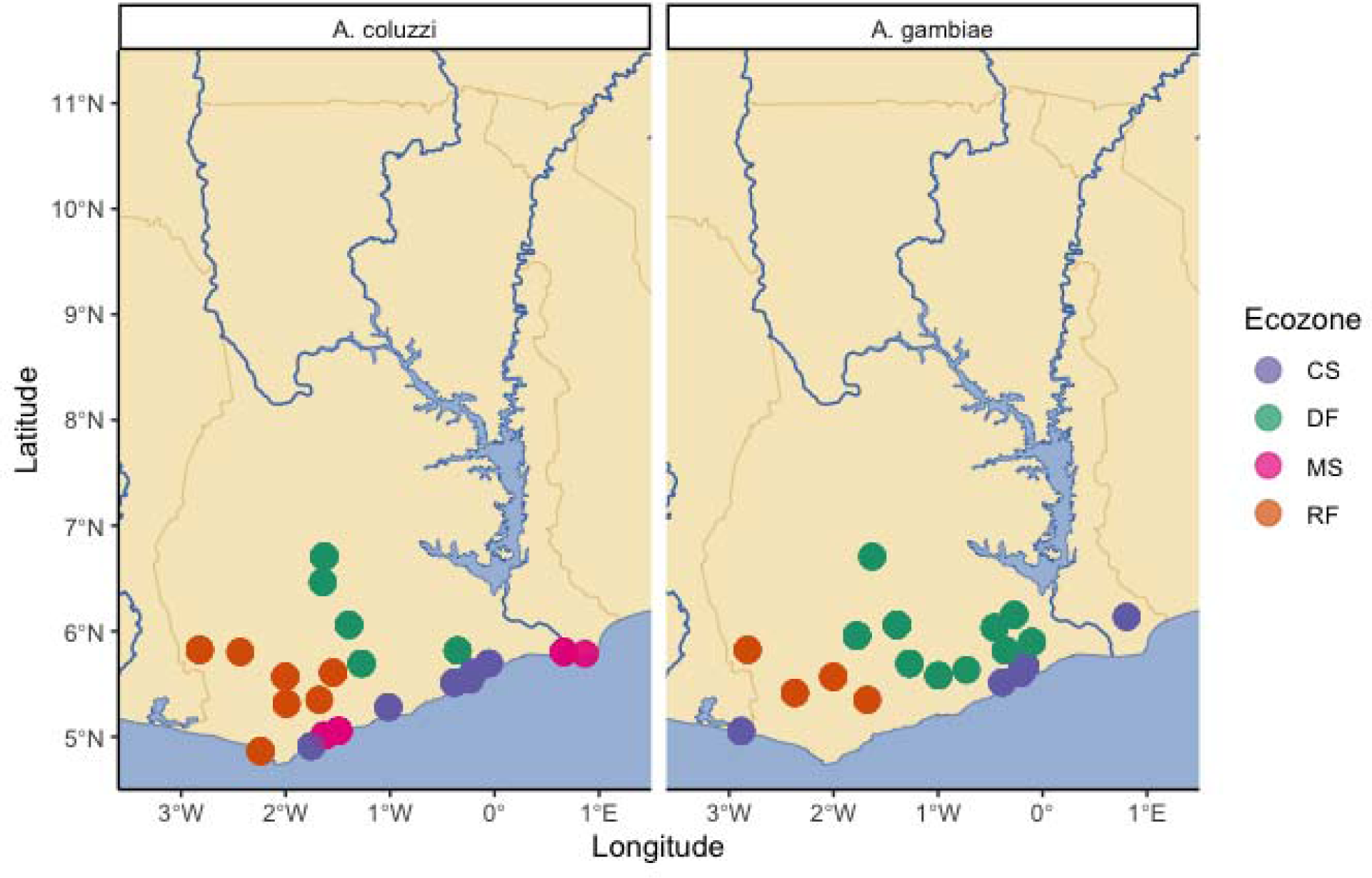
Sample collection scheme for *Anopheles coluzzii* (LH Panel) and *Anopheles gambiae* (RH panel) in Southern Ghana. X axis indicates longitude, y axis indicates latitude, scale bar indicates 200km distance. Point colour stands for ecoregion (CS: coastal savannah, DF: deciduous forest, MS: mangrove swamp and RF: rainforest, respectively).

**Figure 2:**
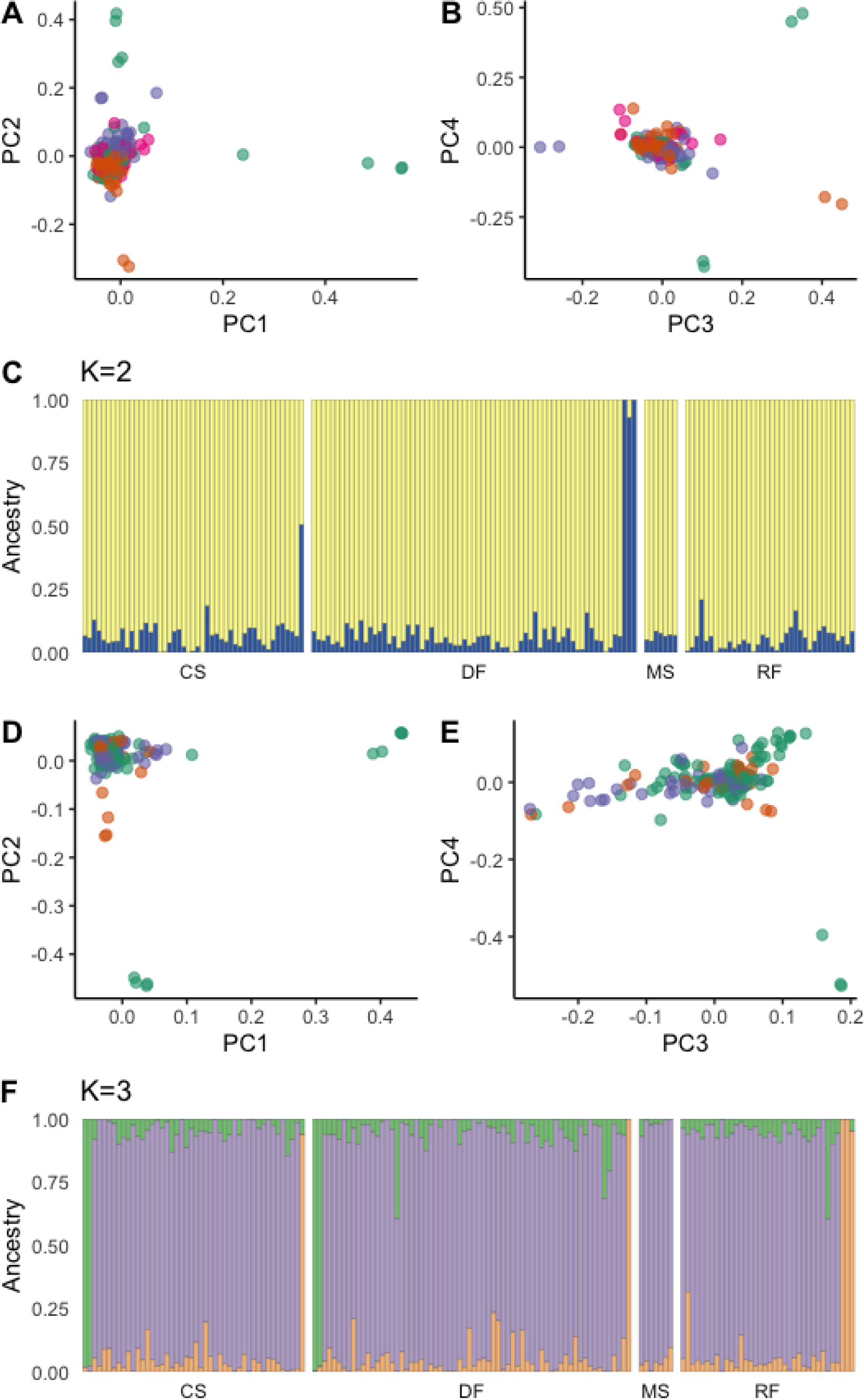
**Panels A-C** denote population structure of *An. coluzzii* in Southern Ghana. Principal component analysis (PCA) plots for principal components (PCs) 1 vs 2 (**A**) and 3 vs 4 (**B**), with points coloured by sample ecoregion. Panel **C** depicts the most likely value of *K* in an ADMIXTURE analysis; the Y axis is the admixture proportion of a given cluster (colour) for a single individual (X axis), from one of the four ecoregions sampled (defined in Fig 1). **Panels D-F** indicate Population structure of *An. gambiae* in Southern Ghana. Principal component analysis (PCA) plots for principal components (PCs) 1 vs 2 (**C**) and 3 vs 4 (**D**), with points coloured by sample ecoregion. Panel **E** depicts the most likely value of *K* in an ADMIXTURE analysis, whereby the Y axis is the admixture proportion of a given cluster (colour) for a single individual (X axis), from one of the four ecoregions sampled.

**Table 1:**
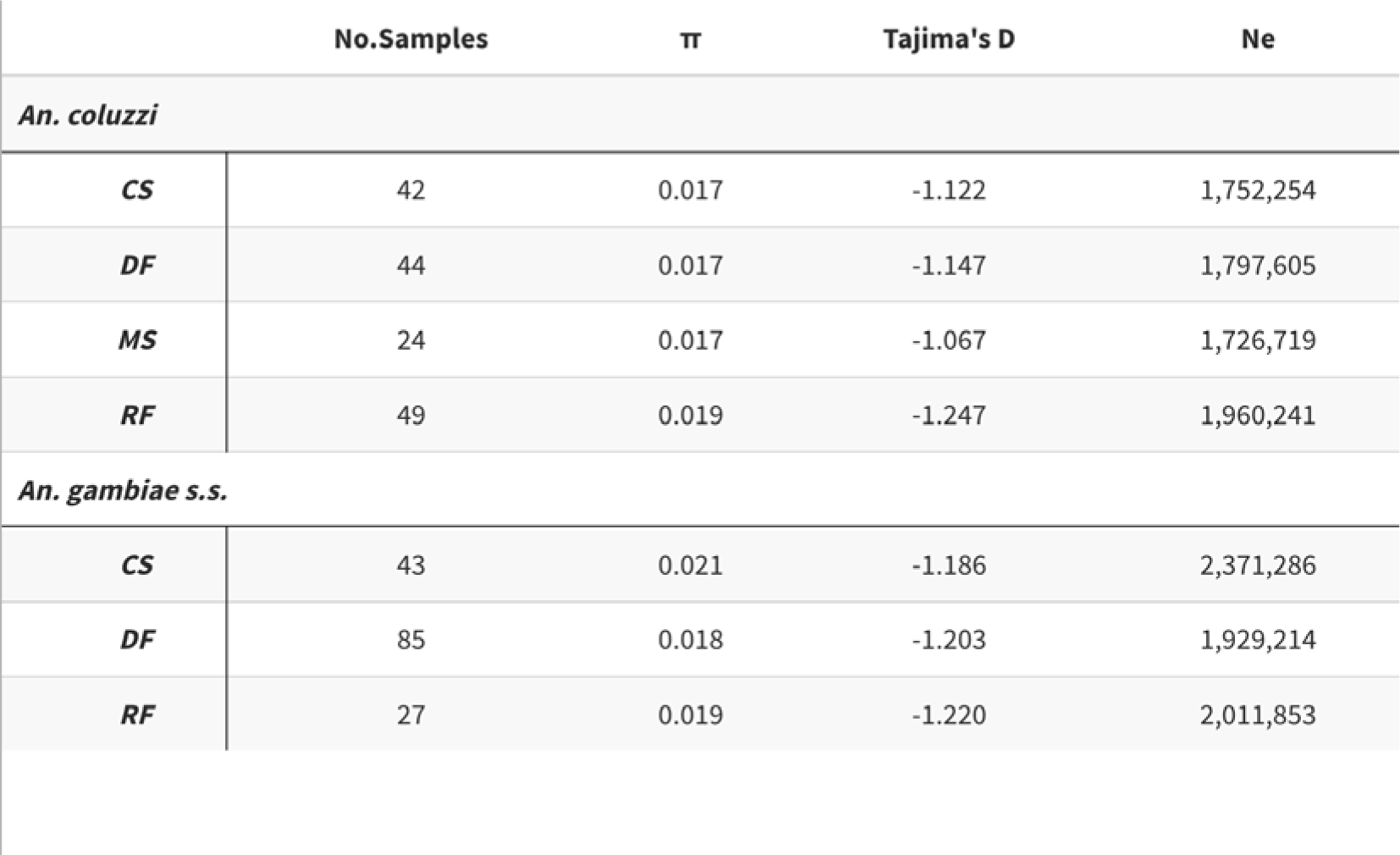
Number of samples sequenced,and genomewide mean values of π, Tajima’s D and *Ne* per species and ecoregion.

### Diversity and differentiation by ecology and species

We characterised genomewide differentiation (*Fst*) within *An. coluzzii* and *An. gambiae* between ecological zones. (**Tables 2A-2B**). Each pairwise *Fst* between *An. gambiae* ecoregions was higher than those in *An. coluzzii* (**Table 2**). In *An. coluzzii, Fst* between forest ecoregions (DF:RF - 0.002) and non-forest ecoregions (CS:MS - 0.002) were lower than between forest and nonforest ecoregions (CS:RF - 0.004, MS:RF - 0.004, CS:DF - 0.003, MS:DF - 0.004) (**Table 2A**). Similarly in *An. gambiae* (though there was only one nonforest ecoregion: CS), *Fst* between forest (RF:DF 0.005) was lower than forest:nonforest (CS:RF - 0.041, CS:DF - 0.008). We calculated the mean genomewide Tajima’s D and π, along with estimated *Ne*, which are shown in **Table 1**. Values of mean π varied between 0.021 (*An. gambiae* DF) and 0.017 (*An. coluzzii* DF), and those for mean Tajima’s D between -1.067 (*An. coluzzii* MS) and -1.247 (*An. coluzzii* RF). Estimates of mean *Ne* varied between approximately 1.7 million (*An coluzzii* MS) and 2.4 million (*An. gambiae* CS). *An. coluzzii* π and *Ne* were consistently lower than *An. gambiae,* and Tajima’s D consistently less negative across comparable ecoregions. (**Table 1**).

**Table 2:** Mean genomewide *Fst* between major ecoregions in southern Ghana for (**A**) *Anopheles coluzzii* and (**B**) *An. gambiae* samples.

| A | CS | DF | MS |
| --- | --- | --- | --- |
| RF | 0.0039 | 0.0023 | 0.0042 |
| CS |  | 0.0027 | 0.0017 |
| DF |  |  | 0.0037 |

| B | CS | DF |
| --- | --- | --- |
| RF | 0.0145 | 0.0054 |
| CS |  | 0.0076 |

We identified a total of 7877 genes located in outlier regions of relatively increased *Fst* (**see Materials and Methods**) between ecoregions in *An. coluzzii* and *An. gambiae* combined (**Supplementary Table 2**). The majority (4827 - 61.27%) of these genes were located as part of a general elevation of *Fst* in the *2La* inversion, making it difficult to identify specific genes therein which are potentially under selection. We identified numerous peaks centred on, or close to, genes implicated in insecticide resistance. Though windows in peaks with the highest *Fst* were not always those containing IR genes, which were sometimes close or adjacent, we considered it likely that the IR genes might typically be targets of selection (See **Materials and Methods**). On *An. gambiae* chromosome 2R, we observed a sharp outlier peak centred on the *Ace1* locus in all 3 ecoregions, particularly striking for the RF comparisons, and at the *Cyp6P* complex of genes, between all 3 ecoregions and highest for CS comparisons (**Figure 3**). We were unable to identify any specific genes underlying the peak between DF:CS at ∼40Mb on chromosome 2R, or in the pericentromeric region between ∼46-60Mb. On *An. gambiae* chromosome 2L, the signal was dominated by the *2La* inversion with the highest *Fst* between CS and RF, suggesting strong contrast in inversion polymorphism frequencies. On 2L, we noted other striking peaks, centred on the *Vgsc* gene at approximately 2Mb, and at genes of unknown function (AGAP005300 and AGAP005787) in the comparisons between forest and non-forest regions. On chromosome 3R an outlier *Fst* peak - at approximately 28Mb - contained a number of genes of unknown function but is located approximately 20kb from the epsilon *Gst* cluster containing *Gste2*. (**Figure 3**). Additional peaks between RF and non-RF zones on chromosome arm 3R are located in the pericentromeric region that contain a range of genes, including potential IR candidates *acyl-coA synthetase* and *Cyp303a1*. Clear peaks were less evident on chromosome arm 3L but on chromosome X, two major peaks were obvious, one centred on the *diacylglycerol kinase (ATP-dependent)* gene at approx 9Mb, and the other outlier *Fst* peak centred on the *Cyp9k1* gene at approx 15Mb (**Figure 3**). This was the highest region of differentiation genomewide for any of the three ecoregion comparisons in *An. gambiae*.

**Figure 3:**
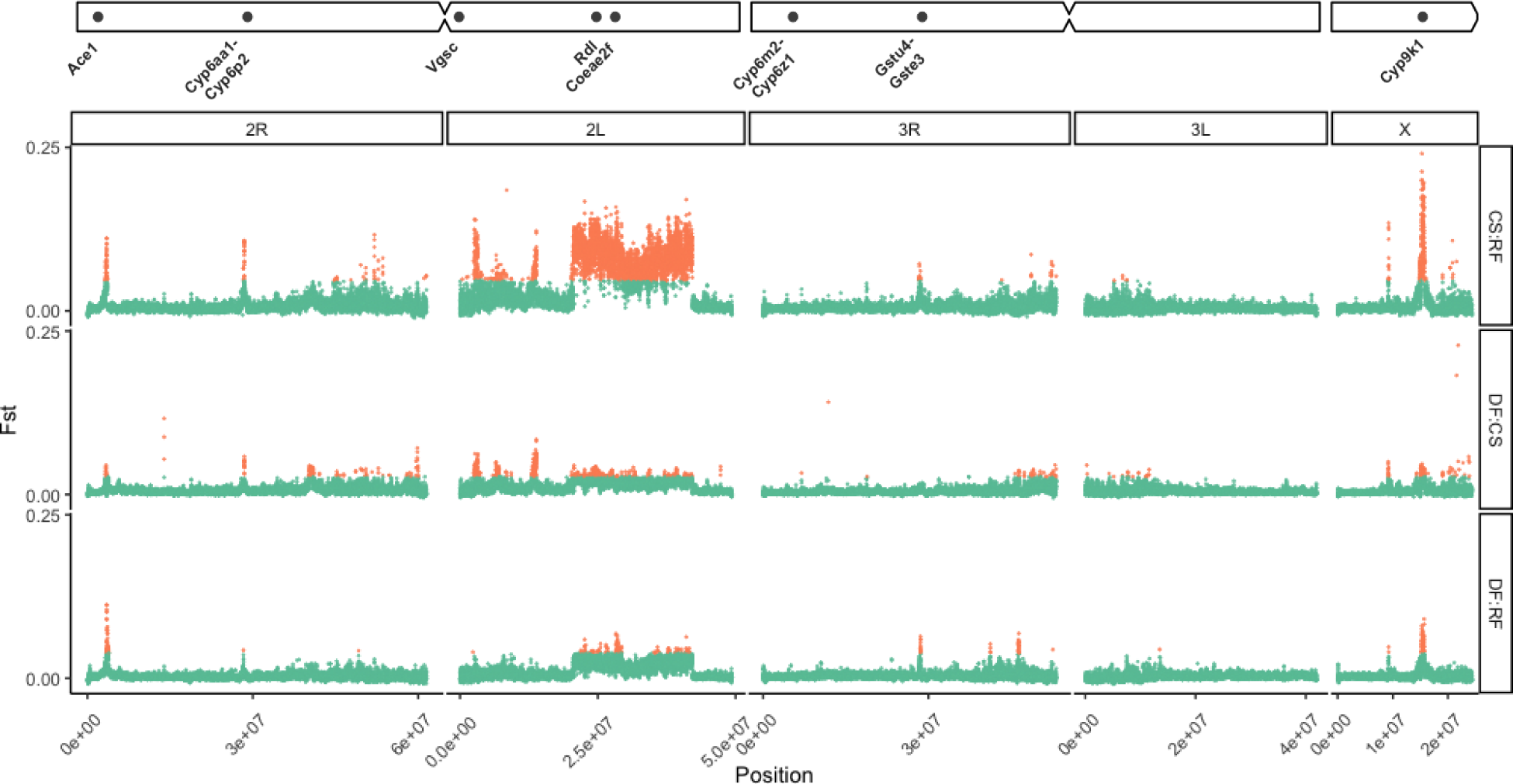
*Fst* genome scan (10kb windows with 5kb step) between *An. gambiae* larvae from deciduous forest (DF), coastal savannah (CS), and rainforest (RF) from southern Ghana. The x-axis indicates chromosomal position (bp). The y-axis indicates *Fst*, orange point colour indicates that a given window was designated as an outlier. A chromosome ideogram with IR gene positions is plotted at the top.

For *Anopheles coluzzii,* (**Figure 4**) there were fewer *Fst* peaks between ecoregions than in *An. gambiae*. Generally, there appeared to be very peaks of differentiation between forest (DF:RF) ecoregions. (**Figure 4**). On chromosome 2R, there were consistent peaks at approx 28Mb around the *Cyp6-*complex containing region, and less consistently a peak at ∼3.4Mb around the *Ace1* locus (**Figure 4**). On chromosome 2L, we found a striking peak on 2L between CS and non-CS zones centred on a window containing the *Pyruvate dehydrogenase E1 component subunit alpha* and *alpha-tocopherol transfer protein-like* genes (**Figure 4**). Against a backdrop of generally higher *Fst* between CS:RF and other ecoregion comparisons, we found relatively small peaks along 2L, including *Vgsc* and, notably, no 2La differentiation. In addition to the *Vgsc* peak, we also saw a small peak between MS and CS at approx ∼38Mb at the AGAP029693 *and amiloride-sensitive sodium channel* genes. There were very few *Fst* outlier peaks on 3R and 3L, with outlier regions mainly concentrated in broad peaks in the pericentromeric regions (**Figure 4**). The most distinct peak on 3R was a peak at approximately 32Mb that contained a number of odorant receptor (Or18-53) genes, as well as a small peak at ∼4.3Mb that contained a complex of *Cyp12F* genes) - this peak appeared between CS and non-CS zones in *An. coluzzii* but was not designated as an outlier in other comparisons outwith CS:RF (**Figure 4**). The most notable *Fst* outlier peaks were consistently between forest (DF/RF) and nonforest (MS/CS) ecoregions at the *Cyp9k1* locus, with much lower peaks between DF and RF or MS and CS.

**Figure 4:**
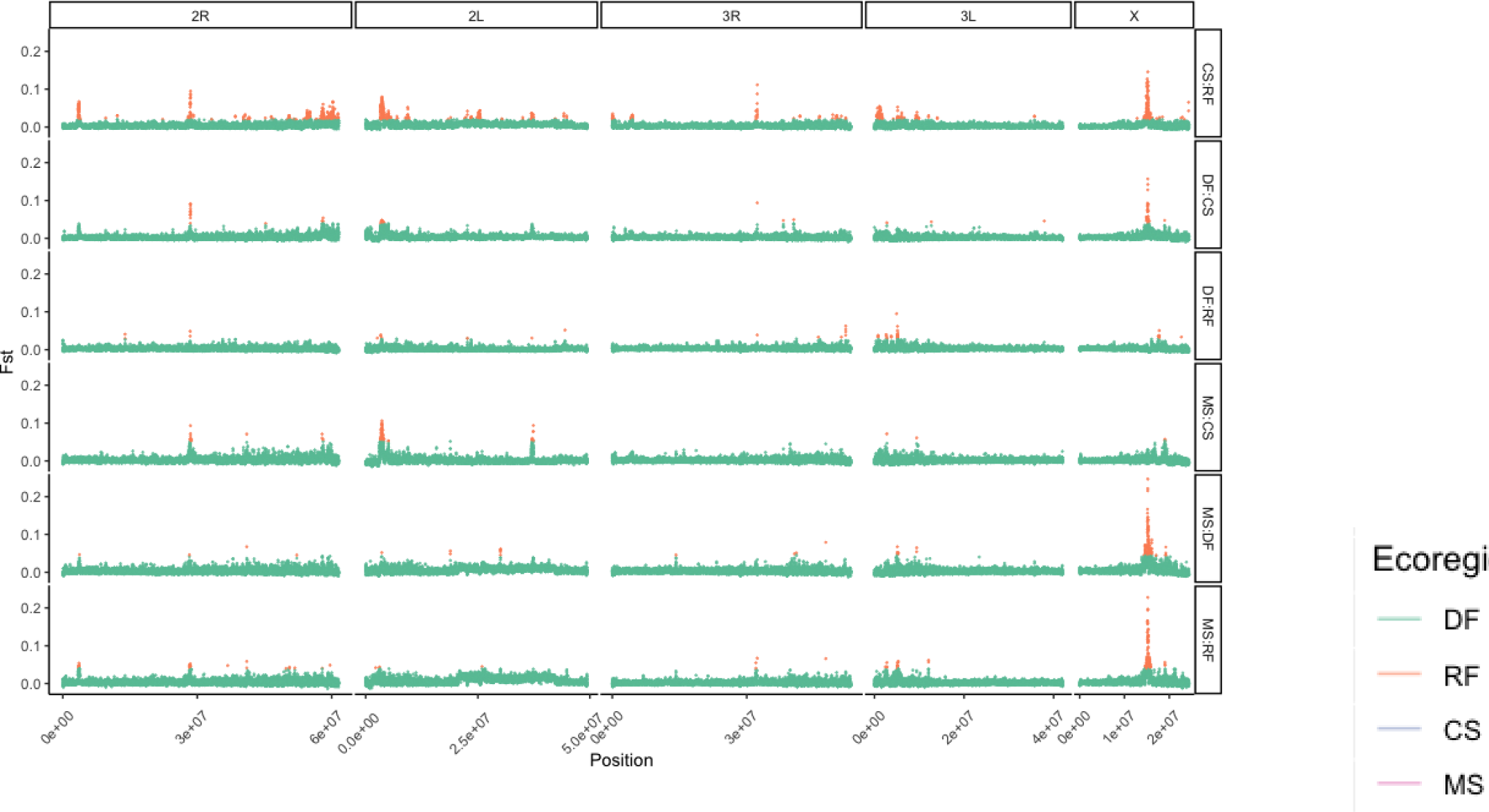
*Fst* genome scan (10kb windows with 5kb step) between *An. coluzzii* larvae from deciduous forest (DF), coastal savannah (CS), mangrove swamp (MS), and rainforest (RF) from southern Ghana. The x-axis indicates chromosomal position (bp). The y-axis indicates *Fst,* orange point colour indicates that a given window was designated as an outlier. A chromosome ideogram with IR gene positions is plotted at the top.

### Selection

*Fst* can identify signatures of selection by looking between populations. We further investigated genomic selection *within* populations by using H123 scans of per-species and per-ecoregion phased GLs (**See Materials and Methods**). In *Anopheles coluzzii*, elevated H123 was apparent in the *Cyp6-*containing region on chromosome arm 2R, at the *Vgsc* and *Rdl* - containing regions on chromosome arm 2L, in the epsilon *Gst* cluster containing *Gste2*, on 3R, and the *Cyp9k1* - containing region on chromosome X (**Figure 5A**). In H123 scans of *An. gambiae,* (**Figure 5B**) notable peaks included the windows containing the *Cyp6* cluster on chromosome 2R, *Vgsc*, *Rdl* and *Coeaef* on chromosome 2L, the *Gst*-epsilon containing region on chromosome 3R, and the *Cyp9k1* and the *diacylglycerol kinase (ATP-dependent)* genes on chromosome X. Whilst signatures of elevated H123 largely recapitulated the *Fst* results from **Figures 3** and **4**, peak sizes were more consistent across the ecoregions, than when comparing between ecoregion types with *Fst*, e.g. at the *Cyp9k1* locus.

**Figure 5:**
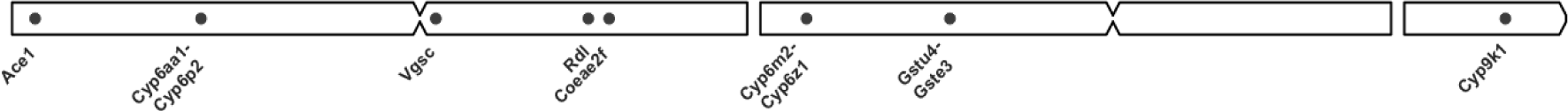
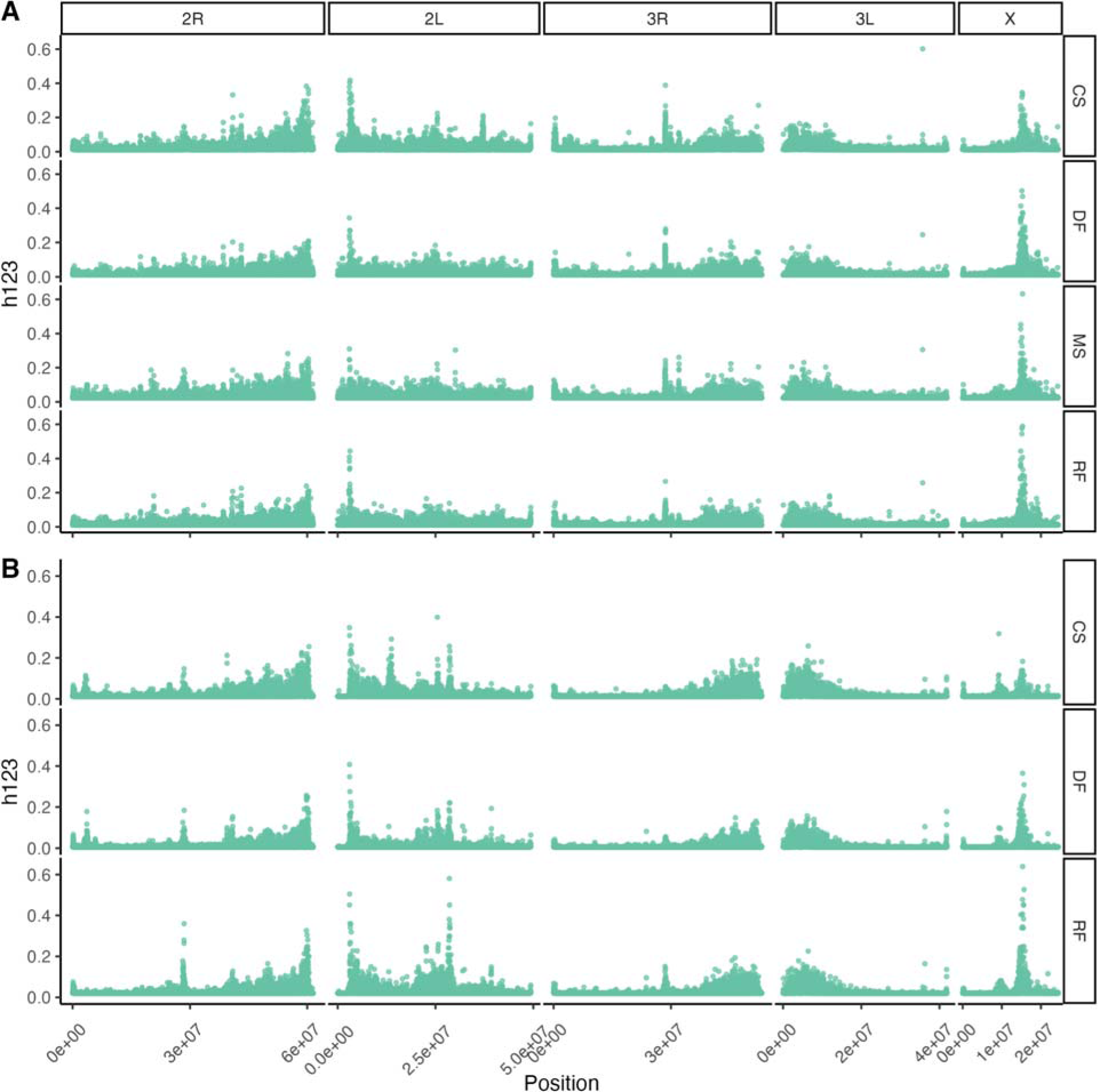
Garud’s H12 scans (in windows of 100 phased GLs) in (**A**) *An. coluzzii* and *An. gambiae* (**B**) larvae from deciduous forest (DF), coastal savannah (CS), mangrove swamp (MS), and rainforest (RF) (row wise panels) from southern Ghana. The x-axis indicates chromosomal position (bp). The y-axis indicates Garud’s H123. A chromosome ideogram with IR gene positions is plotted at the top.

### Variation at the Cyp9k1 locus

The strongest signals of selection were present at the *Cyp9k1* gene. Upon further investigation, we found 19 possible polymorphic sites spanning the locus in *An. coluzzi* (**Table S3**), and 16 in *An. gambiae* (**Table S4**). These were a mix of 3’UTR, 5’UTR, synonymous, intronic variants, with a single missense variant, present only in *An. gambiae* (p.Asn224Ile) (**Table S3**). This variant, p.Asn224Ile was highest in frequency in CS (0.61), followed by DF (0.47), then CS (0.31) (**Table S3**).

### Geographic population structure

We attempted to identify signatures of geographic population structure through the decay in between-sample kinship with geographic distance (isolation-by-distance). (**Figure S1**). The estimated KING kinship coefficient between samples varied between -2.066 and 0.293, with a median of -0.250, for *An. coluzzii,* and -1.067 and 0.370 with a median of -0.265 for *An. gambiae*. We identified 9 full-sib pairs (see **Materials and Methods**) in *An. coluzzii* and 16 full-sib pairs in *An. gambiae*. We identified only one full-sib pair from the same site in *An. coluzzii,* and none in *An. gambiae*. Mantel tests for isolation-by-distance showed no statistically significant signal of isolation-by-distance for *An coluzzi* (*r=*0.009 *p=*0.49) or *An gambiae* (*r=-*0.05 *p=*0.90. In addition, we attempted to identify isolation-by-distance using between-site *Fst* (**Figure 6A**), and found that, like relatedness, Mantel tests for isolation-by-distance were insignificant in *An. coluzzii* (r=0.0008, *p=*0.485) and *An. gambiae* (r=-0.05*, p*=0.898). However, in Mantel correlograms showing correlation between genetic and geographic distance classes (**Figure 6B**), we found evidence of significant isolation-by-distance in *An. coluzzii* at a distance class between 50 and 100km (r=0.14, *p=*0.02), but none in *An. gambiae*. (**Figure 6C**).

**Figure 6.**
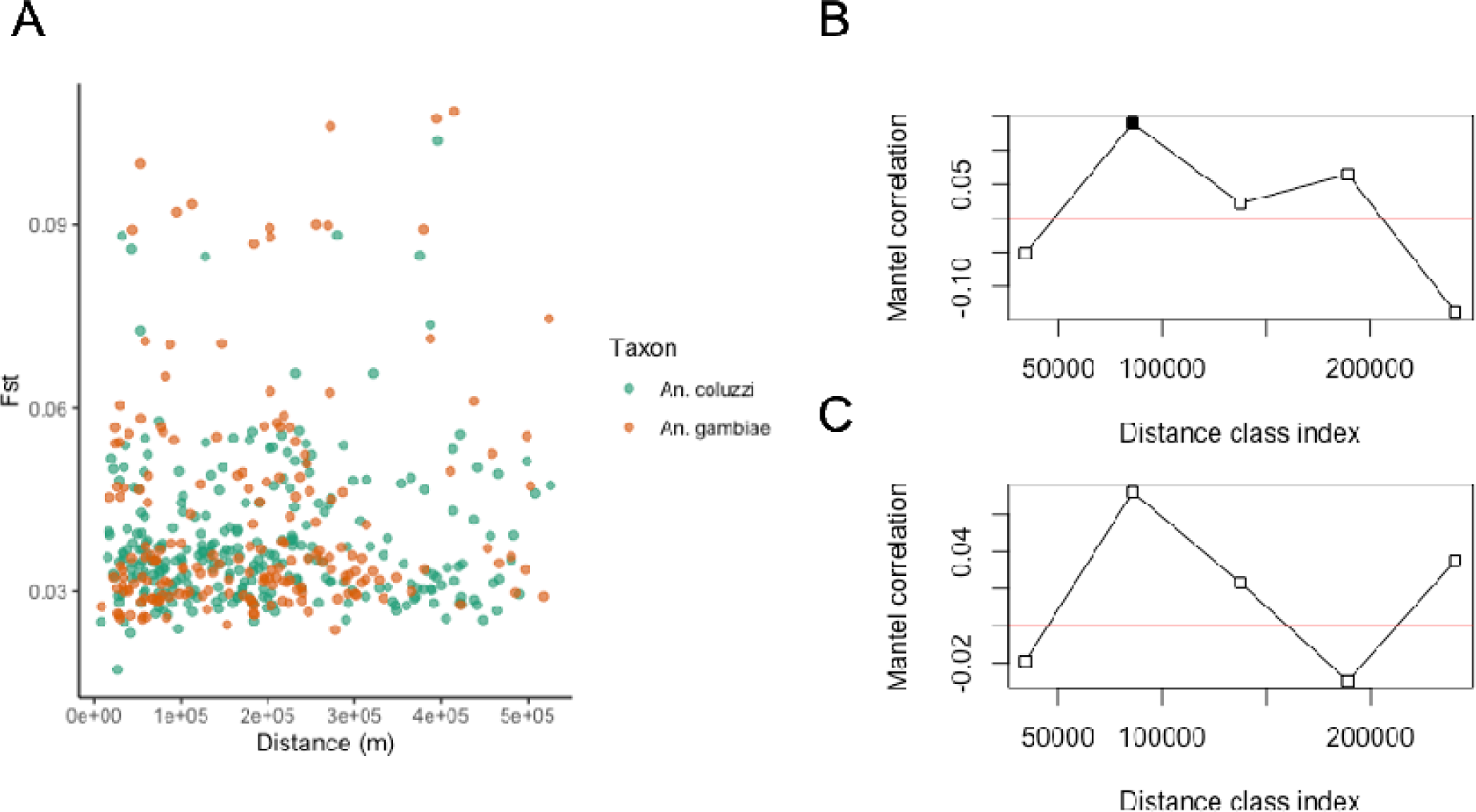
**(A)** *Fst* plotted against geographic distance between sites for *An. gambiae* (blue) and *An. coluzzii* (red). Mantel correlograms showing spatial correlation values over spatial distance classes for *An. coluzzii* (**B**) and *An. gambiae* (**C**).

### Downsampling analysis

We determined the extent to which signatures of population structure, genetic differentiation, relatedness, and selection in *Anopheles gambiae s.l.* may be robust to low sequencing depth-of-coverage (DOC) by performing a set of analyses on a restricted set of samples with a DOC of 10X or over, *in silico* downsampled to 10, 7.5, 5, 2.5, 1 and 0.5X. (**Figure 7**) In relatedness analyses of downsampled individuals (**Figure 7A**), we found that, for pairs of individuals with detectable relatedness, estimates of the kinship coefficient *Rxy* reduced on average with every stepdown of DOC, with the largest reductions being between 5, 2.5 and 1X (N.B. 0.5X was not included in this analysis as kinship for almost all samples reduced to zero at this DOC level). The major grouping patterns in PCA were robust to a DOC as low as 1X, though at 0.5X, the major signals of differentiation appeared to be disrupted. (**Figure 7B**). In genome scans of *Fst* between *An coluzzii* and *An gambiae* chromosome X (**Figure 7C**) *Fst* changed little when DOC was reduced to as low as 1X, whereas at 0.5X, *Fst* appeared to increase in general along the chromosome. In H123 scans incorporating phased genotype-likelihoods, the signal of selection (i.e. elevated H123) on *An. coluzzii* chromosome X appeared similar between coverages of 10X - 7.5X, raised slightly at 5X-2.5X, and lower or absent at 1 and 0.5X (**Figure 7D**). The concordance of downsampled sets of phased *An. coluzzii* GLs appeared to be very sensitive to DOC, with marked increases in discordance rate between the full- and reduced-DOC GL set,with every stepdown of DOC, until 1-0.5X where similarly poor concordance was seen (**Supp Figure 3**).

**Figure 7:**
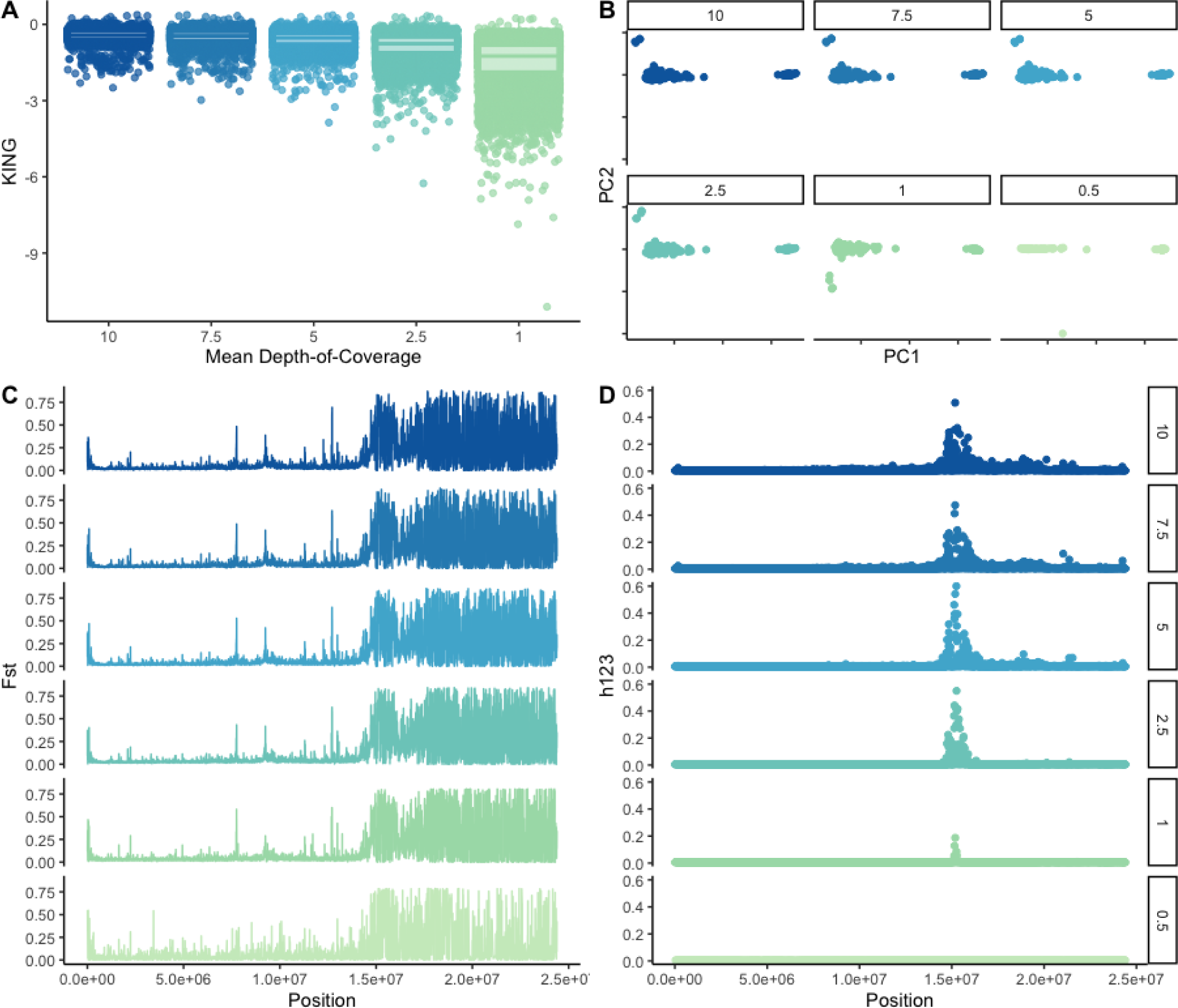
***In silico downsampling results:*** 219 samples > 10X were *in silico* downsampled from 10X to 0.5X (labelled).(**A**) Pairwise kinship coefficients *Rxy,* inferred between 219 pairs of *An gambiae s.l* from Southern Ghana. The y-Axis indicates inferred kinship coefficient; the x-axis indicates *in-silico* downsampled depth-of-coverage.(**B**) PCA results by downsampled coverage **(C)** *Fst* genome scan (2kb windows) of X chromosome between 219 *An. gambiae* and *An. coluzzii* larvae, sequenced to at least 10X, from southern Ghana. The x-axis indicates chromosomal position (bp), with chromosome arm labelled below; the y-axis indicates *Fst*. (**D**) Garud’s *h123* selection scan of the X chromosome in *An. coluzzii* larvae, sequenced to at least 10X, from Southern Ghana. The x-axis indicates genomic position; the y-axis indicates *h123*. Subpanels in **C** and **D** show downsampled DOC (as indicated to the right of **D**).

## Discussion

The results presented in this study suggest that irrespective of ecoregion there are high levels of gene flow and genetic diversity among relatively unstructured populations of *An. gambiae* and *An. coluzzii* across southern Ghana. Isolation by distance was almost absent, with the only significant result coming from comparisons in the 50-100km bracket in *An. coluzzii*, and no relationship between kinship and distance in either species. Selection at IR loci appears ubiquitous across all four ecoregions in both species. Observed values of π and Tajima’s D, as well as an apparent lack of genetic structure at a cross-country-wide spatial scale, are consistent with previous WGS data from West African *Anopheles gambiae* species complex.^1,2^ However, some previous studies indicated much stronger differentiation between ecoregions in the *An. gambiae* species pair - for example between mangrove and non-mangrove in Ghana^33^ and between forest and savannah in *An. coluzzii*^2^ and *An. gambiae*^34^. We find that genomewide differentiation between ecoregions (including between the mangrove swamp ecoregion and other areas) is on the whole low, with differentiation concentrated in specific genomic regions.

We observed numerous signatures of selection localised in specific genomic regions, often linked to genes involved in IR. Most of the outlier *Fst*-associated genes were concentrated in inversion *2La* - a genomic region frequently implicated in environmental adaptation in *Anopheles gambiae s.l.*^10,21^ and in other *Anopheles* taxa^38^. Aside from this, the notable *Fst* outlier regions were detected in comparisons between multiple ecoregions in *An. coluzzii* and included outlier windows in peaks at the *Vgsc* between all ecoregions, the *Cyp9k1* locus - particularly striking between forest and non-forest ecoregions, and the *Cyp6* region on chromosome 2R (that was present between all ecoregions in *An. coluzzii*, but particularly in comparisons involving Coastal Savannah). Peaks in H123 were present around other IR loci (e.g. *Gste2*). In contrast to the ecoregion-specific outlier peaks; at these loci, the H123 scans showed signatures of selection at *Vgsc, Cyp6, Cyp9k1* and *Gst-*epsilon (consistent with other studies in West Africa^15,19,20,41,42^) across all four ecoregions in *An. coluzzii*, suggesting that different alleles of these genes may be selected in different ecoregions. In *Anopheles gambiae*, genomewide differentiation was higher in comparisons involving rainforest, most notably CS: RF, consistent with at least some structure between forest and savannah as suggested in previous analyses^2,9,34^. Interestingly, whilst genomewide mean *Fst* was similar between RF and CS or MS in *An. coluzzii*, genomewide outlier profiles showed many more peaks in both species in the CS: RF comparison in both species suggesting possible commonality of selection pressures. RF:non-RF differentiation was particularly marked at the *2La* inversion in *An. gambiae*, as well as at the *Cyp9k1* and *Ace1* containing regions. The cause of these differences in IR allele frequencies by environment is unknown, but it is noteworthy that the possible selection pressure responsible would likely need to be strong enough to counter the homogenising effect of high gene flow. In the absence of an IRS program, selection on markers responsible for carbamate or organophosphate resistance, notably *Ace1,* may indicate ecoregion- or geographically-specific selection pressures in response to agricultural usage of pesticides. In both species, the most striking signal was an elevation in *Fst* between forest and non-forest ecoregion populations, and in all samples for H123, in the genomic region containing the *Cyp9k1* locus - a gene under selection in West African *Anopheles,* implicated in resistance to pyrethroid insecticides.^2,12,19,22,41,42^, having undergone extensive copy number variation^43^. The data we present here suggest that although differentiation between ecoregions varies at *Cyp9k1* in both species, this region appears to be undergoing a selective sweep in all populations and species. The presence of a missense variant in *An. gambiae,* compared to synonymous and 3’/5’-UTR variants in *An. coluzzii*, suggests that *Cyp9k1* may confer resistance with species-specific mechanisms. The reason for this is unclear. The selection pressure conferred by insecticide resistance is often strong enough for introgression to occur in IR genes (e.g. *Ace1*, *Vgsc*), even in genomic regions highly differentiated between members of the *An. gambiae* species complex^44–46^. The proximity of *Cyp9k1* to the pericentromeric region of the X chromosome, which is often highly differentiated between *An. gambiae* taxa, combined with a mechanism of action that may rely on copy-number variation (and therefore is less dependent on the effect of a single mutation^43^), may account for the species-specificity of this locus.

Further systematic studies incorporating more per-site samples across different ecoregions (the per-site number of samples was generally quite low, between 1-10, making accurate sitewise SFS estimation difficult), will enable identification of specific sites where selection pressures may be acting, and studies employing sequencing modalities that enable the resolution of individual genotypes (as opposed to genotype-likelihoods and allele frequencies) will facilitate genotype-environment associations as well as resolution of specific haplotypes involved in IR at loci that appear to be under selection in this study. Low sequencing coverage in this study (∼10X per-sample) prevented us from calling individual genotypes with confidence ^24,47^. Nevertheless, we found that, consistent with simulation based analyses ^24–26^, sequencing coverages as low as 1X return similar results for PCA and *Fst* (estimated from the SFS). Estimation of kinship, a key metric for assessing vector dispersal distance through close-kin mark-recapture in *Aedes* ^48,49^, is more affected by coverage, though the decrease in *Rxy* between the same pairs of individuals appears relatively slight. However, a lack of robustly called genotypes precluded us from investigating the frequency and environmental association of specific genotypes in regions that appeared to be under selection. The genotype-likelihood phasing we performed in this analysis appeared to be highly sensitive to coverage, suggesting our analysis of individual haplotype effects should be considered suggestive rather than definitive, until further confirmation can be performed. However, the H123 scans appeared relatively more robust to lower coverage than the genotype concordance between downsampled individuals, perhaps because these rely on calculating frequencies of phased GLs as opposed to analysing individual geno- and haplotypes.

The lcWGS approach employed here is promising for future analyses of vector population genomics where individual genotypes are of less interest. Examples of these may include: the identification of vector dispersal distance with close-kin-mark-recapture and other SFS-based approaches such as population demographic history with coalescent modelling^2^; interrogation of changes in diversity, population size and selection during vector control^50^; and exploratory surveys of vector population structure over space and time as a prelude to more in-depth sequencing studies that interrogate genomic regions under apparent selection. This is particularly pertinent given the growing role of genome sequencing in vector surveillance both for discovery of population structure and dynamics, and for control impact and insecticide resistance^2,15,19,51^.

## Materials and Methods

### Sampling and sequencing

Mosquito larvae were collected using dippers from 34 study sites spanning the four major agro-climatic zones in southern Ghana^40^ (**Supp Table 1**, **Figure 1**). The sampled larvae were collected between April 2016 and October 2017 and raised to adults in an insectary. Genomic DNA was extracted individually from each mosquito using Nexttec kits following the manufacturer’s (nexttec™) protocol. Species characterizations into *An. gambiae* complexes were performed using the PCR protocol described by Scott et al., (1993) and further characterized into *An. coluzzii* and *An. gambiae* using protocols described by Santolamazza et al., (2008). The PCR products were visualised under ultraviolet light after electrophoresis using 2% agarose gel stained with Peqgreen dye manufactured by Peqlab Biotechnologie. Eight mosquito samples - of known species - were picked from each site, and in study sites where both *An. coluzzii* and *An. gambiae* were found, 16 samples (comprising eight of each species) were chosen. Genomic DNA was submitted for library preparation and WGS at SNPsaurus, Oregon, USA.

### GL calling and bioinformatics

Data analyses were performed in *R*^52^ [**R Core Team**], incorporating the *ggplot2,* and *geosphere* libraries^53,54^. Read trimming and mapping for all 384 sequenced samples were implemented in *nextflow*^55^. Reads were trimmed with *fastp*^56^, aligned to the AgamP3 PEST reference genome^57^ using *bwa-mem*^58^, and *samtools*^59^ commands *sort, markdup, index*. ANGSD^60^ was used to infer genotype likelihoods (GLs) with the *samtools* genotype likelihood model –*GL 1* and the filters *–maxDepth 6000 –minQ 30 –minInd 0.25*, removing sites with a total depth of > 6000X, a phred-scaled quality score of <30, and sitewise missingness of >0.25, the SNP p-value of > 0.05 (indicating probable lack of polymorphism), and a minor allele frequency of < 0.05. For π and Tajima’s D estimation, the above parameters were used but without filters for polymorphism (i.e. monomorphic sites were included), depth, or minor allele frequency. GL calling and filtering was performed by species and ecoregion for each of the 5 main chromosomes (2L, 2R, 3L, 3R and X). 70 individuals missing more than 25% of genotype-likelihoods were removed from subsequent analyses, leaving 314 individuals from both species.

### Analysis of population structure, differentiation, and diversity

GLs from chromosome 3L were used as input for *pcangsd*^26^, which calculates covariance matrices and most likely individual admixture proportions. The covariance matrix was used for principal component analysis (PCA) with the *eigen* function in R.

Per-subpopulation (e.g. by species and ecoregion) site allele frequencies (saf) were used to calculate folded joint site-frequency spectra (SFS) in *realSFS*^60^, with pairwise comparisons between each species and ecoregion per-species to calculate the *Fst* in 10000bp sliding windows with a 5000bp step. One-dimensional (1d) SFS were calculated from the unfiltered GLs for each species and ecoregion using *realSFS*. Nucleotide diversity (π), theta (for *Ne* estimation), and Tajima’s D were inferred using the *dothetas* option. For the relatedness analysis, whole-genome GLs, (with genomic regions containing inversion polymorphisms masked, as these can lead to spurious estimates of relatedness^19^), were used as input for *NGSRelate*^28^. A pairwise relatedness (KING)^61^ matrix was extracted and compared with a pairwise geographic distance matrix (inferred with the distGeo function from the *geosphere* package in a Mantel test for isolation by distance with *vegan*^62^.

### Identification of outlier genes

We identified genomic windows of potential selection by searching genome-scans of *Fst* for regions of greater than expected differentiation^18^. Outlier windows were designated according to the approach described here^19^, but briefly, for each ecoregion:ecoregion comparison, we took the difference between the smallest *Fst* value, and the modal *Fst,* and designated as outliers any window with an *Fst* more than three times this distance away from the mode on the of the right hand side of the *Fst* distribution. We then intersected outlier windows with the AgamP4 PEST annotation track^57,63^. We reported IR genes located at, or close to, the window of highest *Fst* contained in an outlier peak. Windows with the highest *Fst* in peaks were not always those containing IR genes but, as selection on these genes in response to insecticide pressures is ubiquitous in the region, and occasionally genomic window analysis misses these genes despite their association with resistance phenotypes^19^, we considered it most likely that these are the genes responsible for the peak and reported them as such. A full list of genes is available at **Supplementary Table 2**.

### Selection and haplotype analysis

Genotype-likelihoods for each species and downsampling condition (see ***GL calling*** and ***Downsampling analysis***) were phased using BEAGLE v4^64^. The resulting phased GLs were used to calculate Garud’s H123^65^ in *scikit-allel*^66^. Phased GLs from the region containing the *Cyp9k1* gene on the X chromosome (AgamP4_X:15,240,572 - 15,242,864) were analysed for potential functional effects using *SNPEff v.4.1*^67^, and frequencies of each variant and ecoregion calculated using *scikit-allel*.

### Downsampling analysis

Samples with a mean sequencing depth of coverage (DOC) of > 10 (*n=*214) were randomly subsampled to 10X, 7.5X, 5X, 2.5X and 1X using *samtools view. pcangsd, ngsRelate, and realSFS* were used to generate PCA/admixture proportions, pairwise relatedness between samples, and between-species *Fst* estimates, respectively, in addition to phased GL sets (**see Above**). Genotype concordance between downsampled phased GL sets was calculated using *bcftools stats*^59^.

## Funding statement

J.E. was supported by a Wellcome Trust Training Fellowship to J.E (Grant No. 110236/Z/15/Z). M.V. and T.D. were supported by the European Research Council under the European Union’s Horizon 2020 Research and Innovation Programme, grant no 852957.

## Acknowledgements

We are grateful to Emily Rippon and Dimitra Pipini for laboratory technical support.

## Author Contributions

J.E., A.E.Y and D.W. designed the study, J.E. collected the samples, T.D., J.E. and D.W. analysed the data, T.D., J.E., B.K.M., M.V. and D.W. interpreted the data, T.D. and J.E. wrote the manuscript. M.V., B.K.M and D.W. supervised the project. All authors reviewed the manuscript.

## Data availability

The Docker container and Conda env containing all dependencies and versions for read trimming, mapping, and QC are located at https://github.com/tristanpwdennis/basicwgs. Scripts for GL calling, SFS estimation and all subsequent analyses are available at: https://github.com/tristanpwdennis/td_je_angam_2022. Raw reads are deposited in the European Nucleotide Archive under project accession number PRJEB71887.

## Notes

https://github.com/tristanpwdennis/basicwgs

https://github.com/tristanpwdennis/td_je_angam_2022

### Competing Interest Statement

The authors have declared no competing interest.

