## Supplementary Figures for "Signatures of adaptation at key insecticide resistance loci in *Anopheles gambiae* in Southern Ghana revealed by low-coverage WGS"

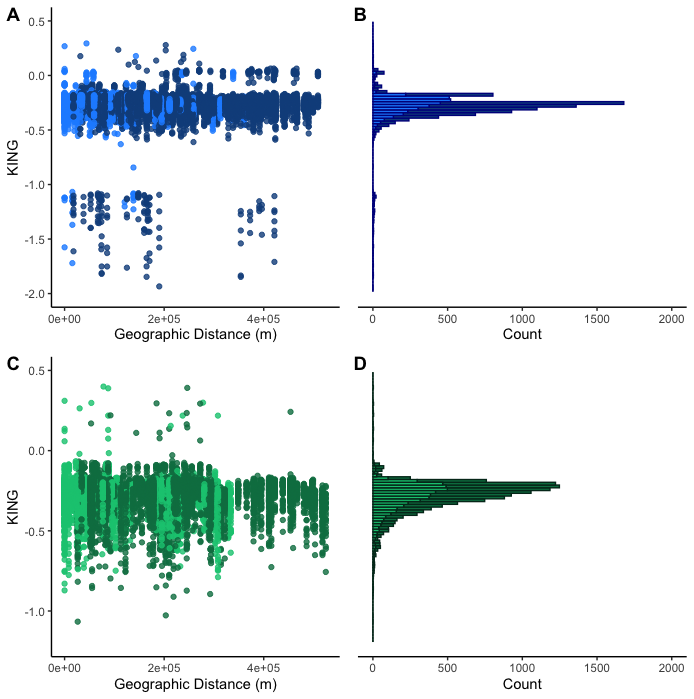

**Supp Figure 1.**Pairwise KING kinship coefficient measured between pairs of *Anopheles coluzzii* (blue, **A,B**) and *An. gambiae* (green, **C,D**) samples. Panels **A** and **C** denote KING coefficient estimates plotted against geographic distance between samples. Panels **B** and **D** are histograms of KING values. Lighter shading indicates samples from the same ecoregion, darker shading, from different ecoregions.

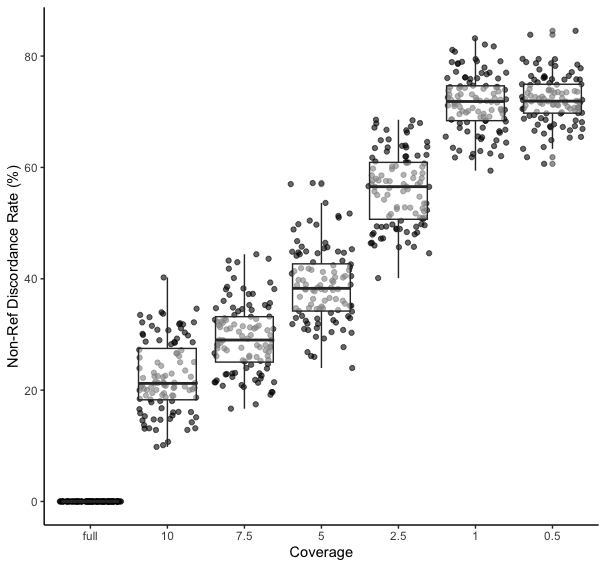

**Supp Figure 2**: Concordance rate (Y axis) of phased genotype likelihoods from *An. coluzzii* at downsampled DOC (X Axis).
